# High-throughput identification of C/D box snoRNA targets with CLIP and RiboMeth-seq

**DOI:** 10.1101/037259

**Authors:** Rafal Gumienny, Dominik J Jedlinski, Georges Martin, Arnau Vina-Villaseca, Mihaela Zavolan

**Affiliations:** Computational and Systems Biology, Biozentrum, University of Basel; Swiss Institute of Bioinformatics

## Abstract

Identification of long and short RNAs, their processing and expression patterns have been greatly facilitated by high-throughput sequencing. Frequently, these RNAs act as guides for ribonucleoprotein complexes that regulate the expression or processing of target RNAs. However, to determine the targets of the many newly discovered regulatory RNAs in high-throughput remains a challenge. To globally assign guide small nucleolar RNAs to site of 2’-O-ribose methylation in human cells, we here developed novel computational methods for the analysis of data that was generated with protocols designed to capture direct small RNA-target interactions and to identify the sites of 2’-O-ribose methylation genome-wide. We thereby determined that many “orphan” snoRNAs appear to guide 2’-O-ribose methylation at sites that are targeted by other snoRNAs and that snoRNAs can be reliably captured in interaction with many mRNAs, in which a subsequent 2’-O-methylation cannot be detected. Our study provides a reliable approach to the comprehensive characterization of snoRNA-target interactions in species beyond those in which these interactions have been traditionally studied and contribute to the rapidly developing field of “epitranscriptomics”.

## INTRODUCTION

RNAs undergo many different modifications in all living organisms (1). High-throughput approaches have been developed recently to map genome-wide 2’-O-ribose methylation (2’-O-Me, (2)) and the most frequent modified nucleobases including N6-methyladenosine (m6A, (3)), pseudouridine (ψ, (4)), and 5-methylcytosine (m5C, (5)). These studies have catalyzed the birth of “epitranscriptomics” (6) and have rekindled the interest in the function of RNA modifications and their relevance for human diseases (7, 8). Whereas 2’-O-ribose methylation has long been implicated in the stability and structure of ribosomal RNAs (reviewed in (9)) and m6A appears to modulate the rate of mRNA translation (10-13), the role of most RNA modifications remains to be characterized.

The 2’-O-methylation of riboses in rRNAs and some other non-coding RNAs such as snRNAs and tRNAs (14, 15) is catalyzed by the protein fibrillarin (16), which is part of a ribonucleoprotein complex additionally containing the 15.5K, NOP56 and NOP58 proteins. The complex is guided to targets by C/D-box small nucleolar RNAs (snoRNA), which take their name from the conserved sequence elements that they contain, known as C/C’ (RUGAUGA, R = A or G) and D/D’ (CUGA) boxes. These are important for snoRNA biogenesis and for interaction with RNA binding proteins (17), A snoRNA interacts with its targets through “anti-sense” elements that are located upstream of the D and/or D’ box. The target nucleotide that pairs with the fifth nucleotide of the snoRNA anti-sense sequence acquires the 2’-O-ribose-methylation mark. Many studies have investigated snoRNA-guided modifications, particularly in yeast (18-21). As a result, some features characterizing functional snoRNA-target site interactions have been inferred and are used in the computational prediction of snoRNA targets (22, 23). For instance, validated snoRNA-target interactions indicate that the 3’ end of the anti-sense box should be highly complementary to the target site, with no more than one mismatch over at least 7 nucleotides and no bulges (23).

Until recently, experimental methods for snoRNA target site identification were laborious and could only address a few sites at a time (24). This has changed with the development of an increased throughput method, CLASH (crosslinking, ligation and sequencing of hybrids), which enables isolation of RNA-RNA hybrids that form between snoRNAs and their targets (25). The approach has also been applied to the identification of microRNA targets (26). Although CLASH has limited efficiency of target capture (27) compared to crosslinking and immunoprecipitation of protein components of ribonucleoprotein complexes (CLIP) (28, 29), it has the advantage that it reveals not only the target site but also the identity of the guide RNA. Interestingly, a recent study found that a small fraction of reads obtained in CLIP experiments represents RNA-target chimeras, which are thought to form due to cellular enzymes ligating the guide miRNA to its target RNA during CLIP (30). Whether the CLIP of core snoRNPs also yields snoRNA-target chimeras has not been investigated.

In parallel to the high throughput capture of snoRNA targets, efforts were undertaken to map all 2’-O-methylated riboses in ribosomal RNAs (2). The novel RiboMeth-seq method takes advantage of the resistance of 2’-O-methylated riboses to alkaline hydrolysis and led to the mapping of 54 annotated and 1 predicted 2’-O-methylated site in S. *cerevisiae.*

Studies from various groups including ours have recently expanded the set of human snoRNAs, beyond those that are cataloged in snoRNAbase (https://www-snorna.biotoul.fr/ (31)) (32-34). Analyzing the small RNA sequencing data sets generated by the ENCODE consortium and taking advantage of the processing pattern that most C/D-box snoRNAs seem to follow (33). we have recently constructed an updated catalog of human snoRNAs (Supplementary Table 1, Jorjani et al. submitted). Many of these molecules identified in small RNA sequencing data as well as snoRNP protein CLIP experiments, contain only a subset of C/D box snoRNA-specific sequence elements and with few exceptions, they lack predicted targets. They are therefore being referred to as snoRNA-like (33). In our previous study (Jorjani et al. submitted) we also found that many snoRNAs are differentially expressed, not only between tissues but also in cancers. This finding suggests that snoRNA-guided 2’-O-ribose methylation plays a role in human diseases.

In the present study we have combined recently published high-throughput experimental protocols with novel computational analysis methods to globally assign guide snoRNAs to 2’-O-Me sites in human cells. We first sought to identify chimeric sequences, composed of a snoRNA and a corresponding target, among the reads obtained by crosslinking and immunoprecipitation (CLIP) of core snoRNPs. To validate the role of CLIP-based identified interactions in 2’-O-methylation we then applied RiboMeth-seq (2). Intersecting high-confidence site sets obtained with the two methods, we have recovered ∽70% of the previously annotated 2’-O-Me sites in human rRNAs. We additionally assigned SNORD30 as guide for the modification at position 1383 of rRNA18S and we found that SNORD2 guides a previously unknown 2’-O-Me modification at position G2435 of rRNA28S. Because the CLIP-based capture of chimeric, guide RNA-target, reads has low efficiency ((30) and our results below), we have also sought supportive evidence for high-confidence 2’-O-Me sites determined with RiboMeth-seq among the low-probability sites inferred from chimeric reads. Additionally, we used the data generated with another protocol that we previously developed to identify modification sites (RiM-seq, see Jorjani et al. submitted). This approach yielded 5 novel sites in rRNAs, two of which were supported also by the RiM-seq data. Some interactions with strong chimeric read support, particularly outside of the canonical snoRNA targets, do not seem to lead to 2’-O-ribose methylation that can be detected with RiboMeth-seq or RiM-seq. This may indicate that C/D box snoRNA interaction with target sites have functions beyond the 2’-O-ribose methylation. Interestingly, these interactions seem to be guided by neurally expressed snoRNAs and involve genes that are associated with neuronal dysfunctions.

## MATERIALS AND METHODS

### CLIP of snoRNP core proteins

To identify chimeric snoRNA-target reads, we analyzed 5 CLIP data sets that were published before (33): 2 NOP58-CLIP (Gene Expression Omnibus accession numbers GSM1067861 and GSM1067862), 1 NOP56-CLIP (GEO accession GSM1067863) and 2 FBL-CLIP (GEO accession GSM1067864 and GSM1067865). We also generated an additional FBL-CLIP data set with the protocol described in (35) (GEO accession # pending).

### Identification of snoRNA-target chimera

#### SnoRNA and target sets

We obtained the most comprehensive annotation of human snoRNA sequences, genome coordinates and known or predicted targets from the human snoRNA atlas generated by (Jorjani et al. submitted). RNAs that are known targets of snoRNAs (rRNA and snRNA) were downloaded from the snoRNA database (31). tRNA sequences were obtained from GtRNAdb (36), and one tRNA sequence per codon was added to the set of putative snoRNA targets. The target database thus consisted of the hg19 version of the human genome assembly, enhanced with rRNA, snRNA and tRNA sequences.

### Computational analysis of chimeric reads

Analogous to a study that mapped miRNA-target interactions based on chimeric reads (30), we developed a method applicable to snoRNAs as follows.

#### Read selection

We carried out an initial annotation of CLIP data sets with the CLIPZ web server (37), which provides as output genome-mapped reads with their respective annotations, as well as the unmapped reads. We used clusters of overlapping mapped reads to extract putative target sites and the unmapped reads longer than 24 nucleotides to search for snoRNA-target chimeras.

#### Detection of snoRNAs fragments in unmapped reads

From each snoRNA we generated all possible sub-sequences of 12 nucleotides in length (“anchors”). When an anchor was found in a chimeric read, we aligned the corresponding snoRNA to the chimeric read with a local alignment algorithm (Smith-Waterman, implemented by swalign python package (https://pvpi.pvthon.org/pvpi/swalign) with parameters: match = 2, mismatch penalty = −5, gap open penalty = −6, gap extension penalty = −4). For each chimeric read we retained only the snoRNA(s) with the best local alignment score. Only ∽20% of the initially unmapped reads in our dataset had such ambiguities. To evaluate the significance of the alignment scores, we applied the same procedure to shuffled reads. As shown in Supplementary Figure 1, for most of the reads the score of the alignment with the snoRNA presumed to be contained in the read was much higher compared to the score of aligning the snoRNA to a shuffled version of the read. Thus, as it appears that some unmapped reads indeed include a snoRNA fragment, we analyzed them further as follows. We called the part of a chimeric read that could be aligned to a snoRNA the “snoRNA fragment” and the rest of the read “putative target fragment”. All reads with a putative target fragment of at least 15 nucleotides were considered candidate chimeras which we analyzed further as described below.

**Figure 1.**
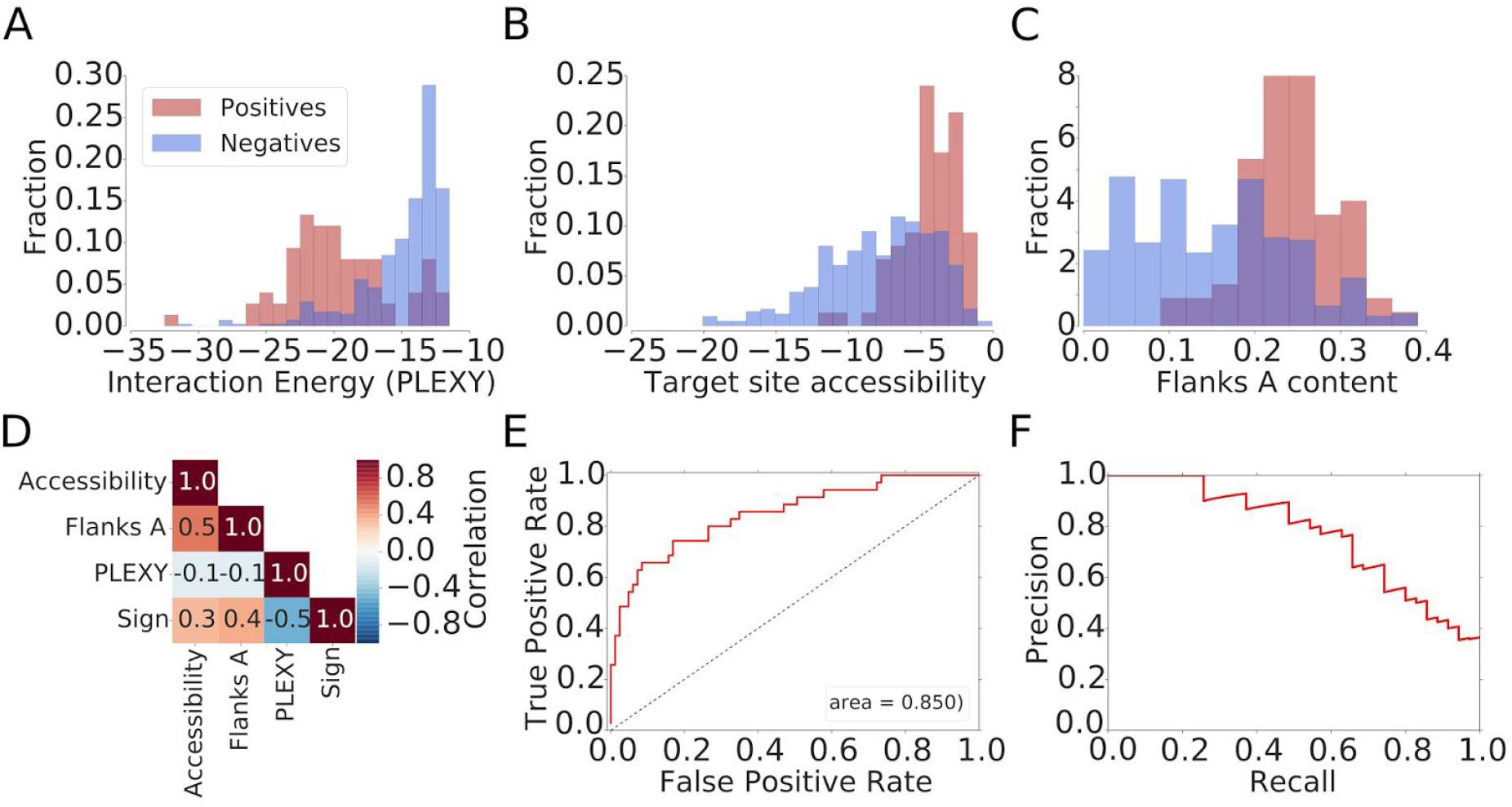
Characterization of the chimeric read-based snoRNA target identification model. Distributions of (A) the interaction energy calculated with PLEXY (22). (B) the target site accessibility calculated with CONTRAfold (39) and (C) the A nucleotide composition of the neighborhood of positive (known) and negative (captured in chimeras but unknown) snoRNA interaction sites. (D) Correlation between features used for model training and the indicator function, taking the value of −1 for negative and 1 for positive sites. (E) Receiver operating characteristic (ROC) curve and (F) Precision-Recall (PR) curve constructed based on snoRNA target predictions in rRNA18S with the model trained on rRNA28S target sites.

#### Annotation of putative target fragments extracted from chimeric reads

Putative target fragments were mapped to CLIPed sites (defined as clusters of at least 2 genome-mapped reads) as well as to rRNA, snRNA and tRNA sequences, which are known to be subject of snoRNA-guided modification. Because the PAR-CLIP protocol yields reads in which C nucleotides are incorporated at the sites of crosslinked U’s, we generated single-point variant of the reads, with one C nucleotide changed to a U before carrying out the mapping of the putative target fragments (30) with Bowtie2 in the local alignment mode. Command line parameters were as follows: -f -D100 -L 13 -i C,1 --score-min C,30 --local -k 10. For reads that mapped to multiple genomic loci, we checked whether at least one of these loci corresponded to a canonical snoRNA target, rRNA or snRNA. If so, we kept only the canonical locus. Otherwise, we kept all putative target loci. The statistics for each experiment can be viewed in Supplementary Table 2.

### Training a model of snoRNA-target interaction

To develop a model that can distinguish *bona fide* snoRNA-target interactions captured in chimeras from erroneously processed reads, we identified putative targets sites that were captured in multiple chimeras with the same snoRNA and had a PLEXY-predicted energy of interaction (22) lower than −12 kcal/mol. In the combined CLIP experiments we identified 67 unique sites known to undergo 2’O-ribose methylation and 295 sites where a modification is not so far known to occur in the 28S and 18S ribosomal rRNAs. For each site we calculated the features described below and trained a model to predict modification sites in the 28S rRNA. We evaluated the performance of the model using the the known modification sites on the 18S rRNA as true positives and all other sites in the 18S rRNA as true negatives. As the performance was high, we combined the two data sets and retrained a model for the comprehensive identification of snoRNA-target interactions.

### Feature definition and computation

#### Interaction energy (PLEXY)

PLEXY is a tool for computational prediction of C/D box snoRNA targets genome-wide, which uses nearest-neighbor energy parameters to compute thermodynamically stable C/D-box snoRNA - target RNA interactions (22, 38) and further applies a set of rules to filter out likely false positives. For each putative target fragment that mapped to the database of putative targets (see section SnoRNA and target sets) we extracted a 50 nucleotides long sequence centered on the target part of the chimeric read, and calculated its interaction energy with the snoRNA in the chimeric read. PLEXY also assigns the position of the snoRNA-induced modification and we kept this information for further analyses. As control, we shuffled the snoRNA associated with each target part in a chimeric reads and repeated the calculation.

#### Target site accessibility

We defined the accessibility of the target region as the probability that a 21 nts-long anchored at the 5’ end at the 7-nts long minimal snoRNA interaction site is in single stranded conformation within an extended region of 30 nucleotides upstream and 30 nucleotides downstream of the minimal interaction site. We computed this parameter with CONTRAfold (39).

#### Nucleotide content of flanking regions

We defined the ‘Flanks A content’ as the proportion of adenines within 30 nt upstream and 30 nt downstream of the 5’-anchored minimal 7-nucleotide long target region predicted to interact with the anti-sense box of the snoRNA. We similarly computed the frequencies of the other nucleotides. Because the frequency of adenines was most predictive of positive interaction sites (Supplementary Figure 2) we only used this feature in the model.

**Figure 2.**
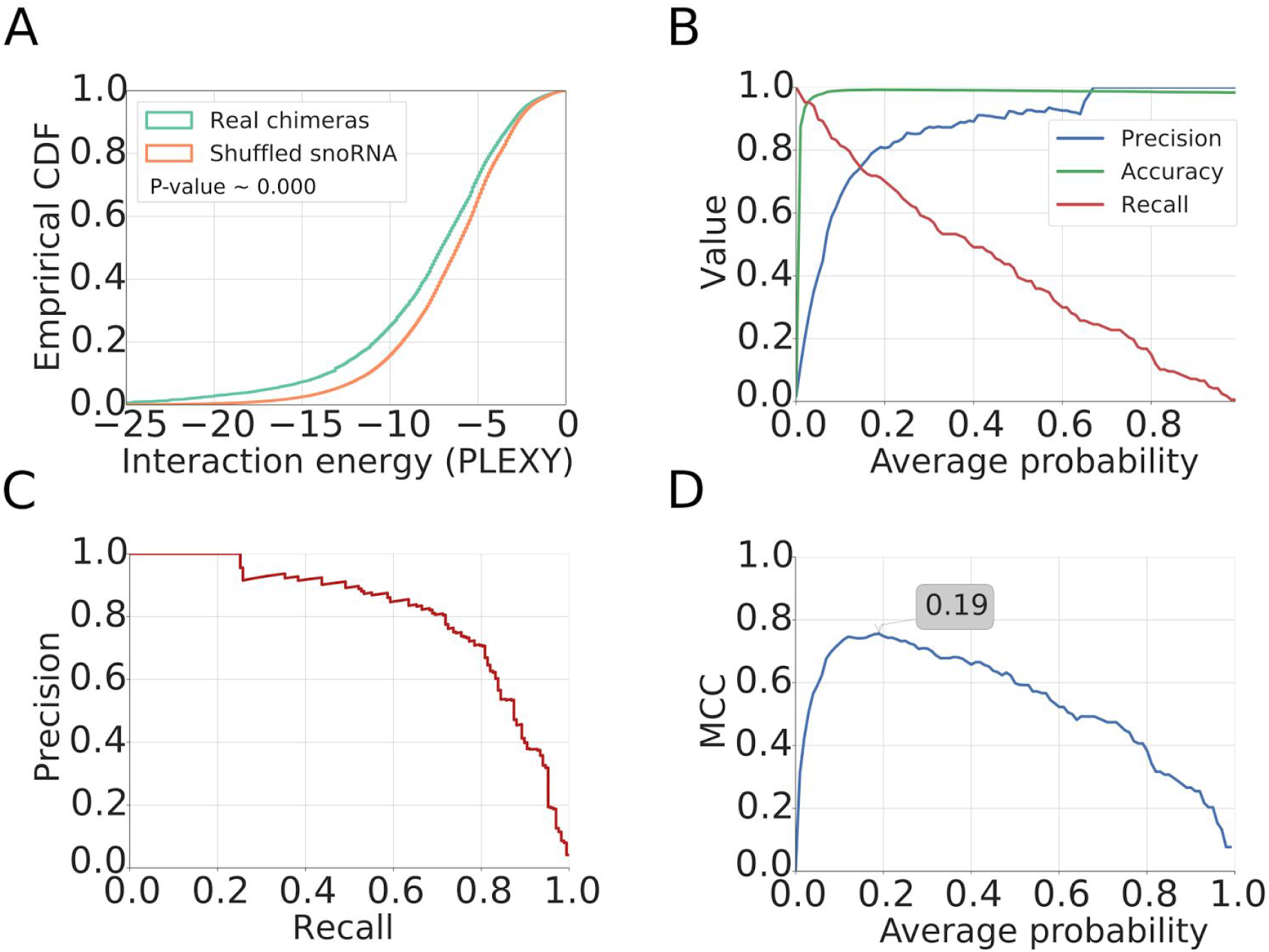
Development of a model for the inference of snoRNA-target interactions from chimeric reads. (A) Empirical cumulative distribution function of the interaction energy estimated with PLEXY between target fragment and snoRNA found in the chimera (Real chimeras) or between target fragment and a randomly assigned snoRNA (Shuffled snoRNA). (B) Metrics illustrating the performance of the method, as a function of the minimum average probability of the considered sites from the 18S and 28S rRNAs across CLIP data sets. (C) Precision-Recall curve for the method. (D) Matthews correlation coefficient (MCC) as a function of the minimum average probability of the considered sites and the derived optimal threshold.

### Model training

The histograms constructed separately for the positive and negative sites in the 28S and 18S rRNAs indicated clear differences between positive and negative sites in the features described above (Figure 1). We used the Statsmodels python library (40) to train a generalized linear model (GLM) with the logit link function (logistic regression). For the comprehensive analysis of the chimeric read data, we used a model which we built based on all rRNA18S and rRNA28S sites (Supplementary Table 3).

#### Annotation of modification sites

We annotated the modification sites based on the ENSEMBL version 75 (41) and the RMSK table from University of California Santa Cruz genome browser (42), for the repeat elements.

### RiboMeth-seq

#### Preparation and sequencing of RiboMeth-seq libraries

To globally map 2’-O-Me sites we adapted the recently developed RiboMeth-seq high-throughput experimental method (2) and applied it to mammalian cells. The principle of the method is that nucleotides with a 2’-O-Me ribose are resistant to alkaline hydrolysis. Thus, products of partial alkaline hydrolysis should not start or end at 2’-O-Me sites, leading to an underrepresentation of these positions among read starts/ends. The read starts/ends will thus provide a negative image of the methylation landscape (2). We applied RiboMeth-seq to map 2’-O-Me sites in rRNA and poly(A)-enriched RNA obtained from either the nucleus or cytoplasmic fraction of HEK293 cells. We also carried out the alkaline hydrolysis for different time intervals of 8, 14 or 20 minutes. The samples that we prepared were as follows:

> RiboMethSeq_HEK_totalRNA_8min
>
> RiboMethSeq_HEK_totalRNA_14min
>
> RiboMethSeq_HEK_totalRNA_20min
>
> RiboMethSeq_HEK_polyARNA_8min
>
> RibomethSeq_HEK_cytoplasmic1_14min
>
> RibomethSeq_HEK_cytoplasmic2_14min
>
> RibomethSeq_HEK_nuclear1_14min
>
> RibomethSeq_HEK_nuclear2_14min

Total RNA was extracted with TRI Reagent (Sigma) and mRNA was prepared with the Dynabeads mRNA DIRECT Kit (Life Technologies), from HEK293 cells according to the manufacturer’s instructions. For mapping of 2’-O-methyl sites in rRNA 1 *μ*g of total RNA was used as starting material. To explore the existence of 2’-O-methyl sites in mRNAs, poly(A)-selected RNA (200ng) was used. In both protocols, the RNA was degraded under alkaline conditions in a sodium carbonate/bicarbonate buffer at pH 9.2 for 14 minutes and then put on ice. Samples were separated parallel to a low molecular weight marker ladder (10-100nt) on a 15% denaturing polyacrylamide gel for 1 hour at 1400 V and 20 W. The gel was stained with GR Green nucleic acid stain (Excellgen) for 3 min and fragmented RNA ranging from 20 to 40 nt was cut out from the gel and extracted overnight in 0.4 M NaCl. The RNA was precipitated with 1 *μ*l of co-precipitant (GlycoBlue) in 75% ethanol at −20°C for 2 hours and then centrifuged at maximum speed for 10 min at 4°C. The RNA pellet was washed twice with 70% ethanol and air-dried. The pellet was dissolved in water, the RNA was dephosphorylated with FastAP alkaline phosphatase (Thermo Scientific) at 37°C for 30 min and the enzyme was heat-inactivated at 75°C for 10 min. Subsequently, the RNA was phosphorylated with polynucleotide kinase (Thermo Scientific) in the presence of 1 mM ATP at 37°C for one hour and then extracted with phenol-chloroform and precipitated in 80% ethanol, washed with 70% ethanol twice and air-dried. The pellet was dissolved in 8 *μ*l mix (4 *μ*l H_2_O, 1 *μ*l 10x truncated T4 RNA Ligase 2 buffer, 1 *μ*l 100 uM 3’ rApp-adapter (5’ adenylated 3’ adapter, 5’-App-TGGAATTCTCG GGTGCCAAGG-amino-3’), 2 *μ*l 50% DMSO), denatured at 90°C for 30 seconds and chilled on ice. Next, RNasin Plus RNase inhibitor (Promega) and T4 RNA Ligase 2 truncated were added to a final concentration of 2 U/*μ*l and 30 U/*μ*l, respectively, and the reaction was incubated at 4°C for 20 hours over night. The next day, 1 μ! of RT primer (100 *μ*M; 5’-GCCTTGGCAC CCAGAGAATTCCA-3’) was added (for quenching of remaining 3’ adapter molecules, preventing adapter dimers ligation in the next step), the samples were heated at 90°C for 30 seconds, at 65°C for 5 minutes, then placed on ice. A 5’-adapter ligation mix was then directly added to the sample (1.5 μ! 10 mM ATP, 1*μ*l 100 uM 5’ RNA Adapter RA5 (Illumina TruSeq RNA sample prep kit), 1 *μ*l T4 RNA Ligase 1 (20 U/*μ*l), 0.5 *μ*l RNasin Plus RNase inhibitor (40 U/*μ*l) and reactions were incubated at 20°C for 1 h and 37° C for 30 minutes. The RNA was then directly reverse transcribed in a 30 *μ*l reaction by adding dNTPs to 0.5 mM, DTT to 5 mM, 1x SSIV buffer, RNAsin to 2 U/*μ*l and 1 *μ*l Superscript IV reverse transcriptase (Life Technologies). The sample was incubated at 50°C for 30 min and inactivated at 80°C for 10 min. 5 *μ*l of the resulting cDNA was then used in a pilot polymerase chain reaction (PCR) reaction. To this end, aliquots were taken from reactions at every second cycle between 12 and 22 cycles and analyzed on a 2.5 % agarose gel. The number of cycles causing a first visible amplification was chosen for a large scale PCR (10 *μ*l cDNA in a 100 *μ*l reaction). The PCR product was ethanol precipitated and run along a 20 bp marker on a 9% non-denaturing polyacrylamide gel in TBE for 1 hour at 250 V, 20 W. The gel was dismantled and stained for three minutes with GR Green. PCR products between 125 bp and 175 bp were cut out, the gel piece was mashed and DNA was eluted overnight into 400 *μ*l of H_2_O. The supernatant was separated from the gel particles in a SpinX filter column (Costar), NaCl was added to 0.4 M, DNA was ethanol precipitated, the pellet washed in 75% ethanol and dissolved in 20 *μ*l H_2_O. Libraries were sequenced on an Illumina HiSeq-2500 deep sequencer (GEO accession # for the data sets pending). Their summary can be found in Supplementary Table 4.

#### Mapping of RiboMeth-seq reads

We obtained ∽50 million reads for each of the RiboMeth-seq samples. The adaptor was removed with Cutadapt (--minimum-length 15, other parameters left with default values) (43) and reads were mapped with STAR (additional parameters: --outFilterMultimapNmax 20 --outFilterMismatchNoverLmax 0.05--scoreGenomicLengthLog2scale 0 --outSAMattributes All) (44) to a human hg19 assembly version-based transcriptome composed of rRNAs, snRNAs and snoRNAs as well as to lincRNAs, miscRNAs, and all unspliced protein coding genes (obtained from hg19 version of ENSEMBL, http://grch37.ensembl.org/index.html) (41), supplemented with a set of tRNAs (as decribed in section SnoRNA and target sets).

#### Calculation of a RiboMeth-seq score

RiboMeth-seq data was evaluated based on “score A”, defined in a previous publication (2) and computed from the mean and standard deviation of the summed coverage of 5’ and 3’ ends at positions neighboring a position of interest. Finding that this score is very sensitive to the coverage and that it does not distinguish very well the true positives (Supplementary Figure 3C), we explored other approaches for analyzing the data. In particular, we tested whether the *change* in cleavage frequency at adjacent positions is indicative of 2’-O-methylation, as we expect very little read coverage at 2’-O-Me positions compared to their immediate neighborhood. Thus, for each target of interest such as the 18S rRNA, we calculated the log2 normalized (to a total library size of 10^6^ reads) profile of cleavage positions from the 5’ and 3’ read ends (separately) and then computed the angle defined by the log2 coverage values at positions −1,0, and +1 with respect to any position along the RNA (Supplementary Figure 3A). An angle of 180° indicates that the three positions have the same cleavage frequencies, 0° indicates that the central position has very high coverage compared to the neighboring positions (and is therefore not protected from cleavage) and 360° indicates that the central position has very little coverage (and therefore it is protected from cleavage) compared to the neighboring positions. Thus, as a RiboMeth-seq score we took the average angle computed based on 5’ and from 3’ read ends, as both ends are determined by alkaline hydrolysis. Calculating the precision, accuracy, recall and precision-recall curves, we found that this score yields a higher precision compared to the ‘score A’ (2) (Supplementary Figure 3B and C). For each individual experiment in predicting 2’-O-Me sites we used a score threshold of 290°, favoring slightly recall over precision. Detail statistics for experiments can be found in Supplementary Table 4. Finally, we combined all RiboMeth-seq experiments and calculated the average score over 7 experiments using the following procedure. We collected putative 2’-O-Me sites from all experiments and retained those that had a score above the threshold in at least one experiment. For these, we calculated the average score across the 7 experiments. To determine a threshold for this average score and then determine the PR curve and Matthews correlation coefficient (Supplementary Figure 3D and E), we included among the positives not only the sites that we identified with RiboMeth-seq but also 19 additional sites that are known to undergo methylation, but that were not captured as positives from our experiments. This resulted in a set of 105 known interactions in the 18S and 28S rRNAs. We found the optimal threshold for the average angle score to be very close to the score for a single experiment, namely 295, giving a recall of 72.3% and precision 84.4% on rRNAs. This indicates that the known 2’-O-methylation sites are captured with very high probability in all RiboMeth-seq experiments.

**Figure 3.**
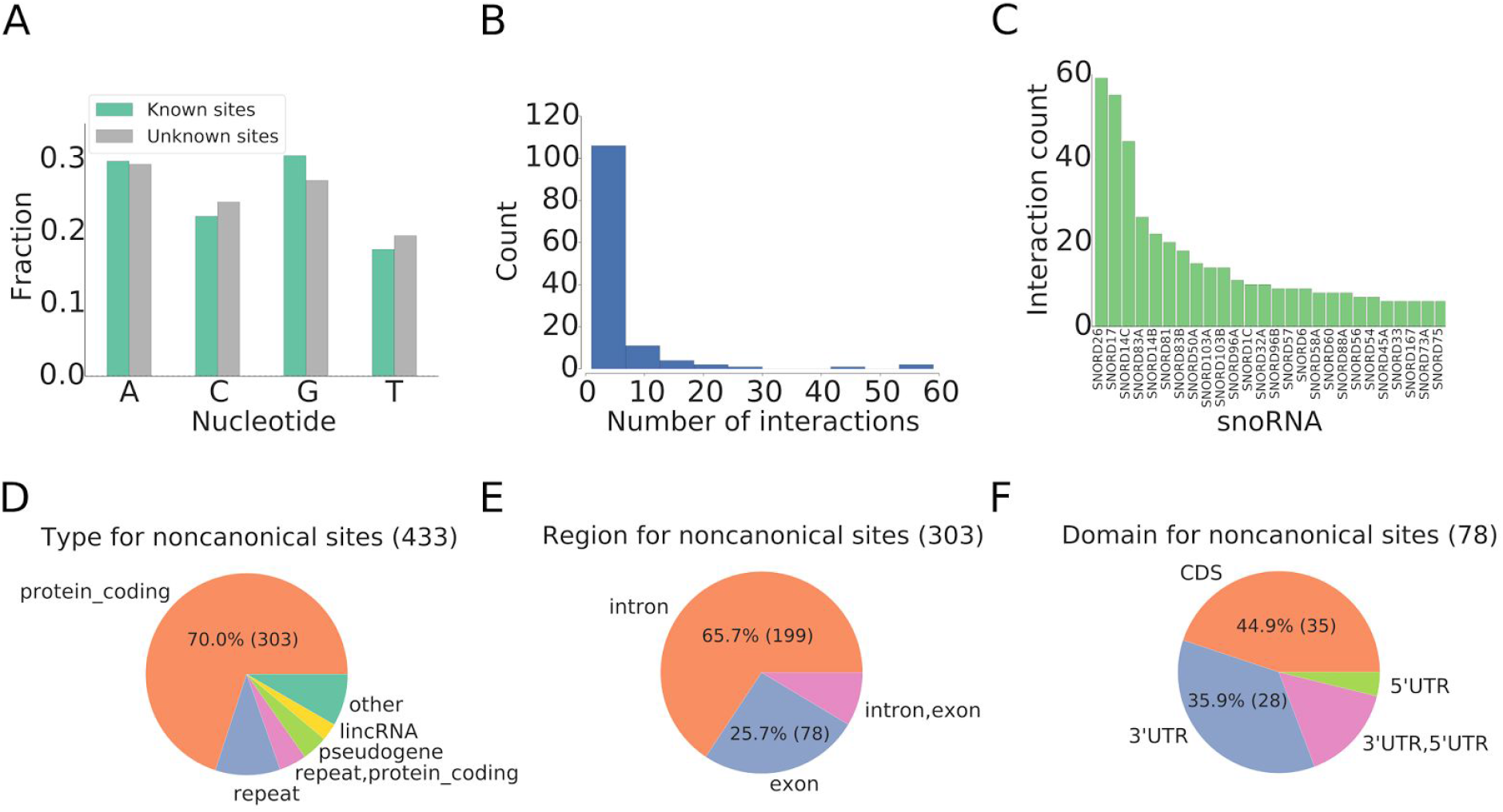
Characterization of novel snoRNA interaction sites observed in chimeric reads. (A) Frequency of RiboMeth-seq-determined 2’-O-Me at the four nucleotides in known and newly identified sites. (B) Histogram of the number of distinct interaction sites identified per snoRNA. (C) Number of distinct interactions of individual snoRNAs. (D) Location of novel interaction sites within transcript types. (E) Relative location of novel interaction sites within protein-coding pre-mRNAs. (F) Relative location of novel interaction sites with respect to mRNA regions.

### Validation of 2’-O-methylation sites with RTL-P

Similar to the classic primer extension assays (45), the Reverse Transcription at Low deoxy-ribonucleoside triphosphate (dNTP) method (RTL-P, (46)) takes advantage of the observation that cDNA synthesis through a 2’-O-Me nucleotide is impaired when dNTPs are limiting. However, RTL-P is less cumbersome and more sensitive than primer extension assays. The RTL-P procedure includes a site-specific primer extension by reverse transcriptase at a low dNTP concentration and a semi-quantitative PCR amplification step, followed by agarose gel electrophoresis to obtain ratios of PCR signal intensities. To increase sensitivity and reproducibility, we implemented a real-time PCR (qPCR) step to facilitate the analysis of the signal intensities (primers shown in Supplementary Table 5).

## RESULTS

### Crosslinking and immunoprécipitation of core snoRNPs captures snoRNA-target site chimeras

Although miRNAs and snoRNAs differ substantially in their mode of action, they share the function of guiding ribonucleoprotein complexes to target RNAs. Thus, by analogy with miRNAs (30), we hypothesized that chimeric reads composed of snoRNAs and their targets should also be captured in CLIP experiments of core components of the snoRNP complex, the 15.5K, Nop58, Nop56 and fibrillarin proteins. Therefore, we analyzed the reads obtained in 6 photoreactive nucleoside-enhanced (PAR)-CLIP experiments that targeted one of these core proteins with a method similar to that previously employed by Grosswendt et al. (30). We found that on average ∽10% of the reads that could not be mapped to the genome or transcriptome seem to be chimeric, composed of a snoRNA and a target part and that ∽5% have the target part longer than 15 nucleotides. Furthermore, in the vast majority of the chimeric reads (∽80%) the snoRNA could be identified unambiguously. The statistics of the reads obtained in all of these experiments are shown in Supplementary Table 2. To determine whether the snoRNA-target associations that we inferred from chimeric reads were due to a *bona fide* interaction between the molecules, we randomized the snoRNA assigned to each target fragment in the chimeras and calculated the predicted energies of interaction of the real and randomized pairs of molecules with PLEXY (22) (see also Methods). As shown in Figure 2A, the interaction energy predicted for the snoRNA-target chimeras was significantly lower compared to randomized sequence pairs, indicating that the chimeras reflected specific interactions of snoRNAs with their targets.

### A model to identify high-confidence snoRNA-target chimeras

Reasoning that features beyond the predicted energy of interaction between a snoRNA and a putative target site may also be informative in identifying true modification sites, we divided the interactions that were captured in chimeric reads into known (positives) and yet-unknown (likely negatives) and compared the distributions of values of features that have been found to play a role in other small RNA-guided interactions (47). As shown in Figure 1A and D, the PLEXY score (22) of the putative snoRNA-target interaction had the highest discrimination power. In addition, we found that snoRNA-guided methylation sites seem to reside in structurally accessible (single-stranded) regions of rRNAs (Figure 1B) and in an A-rich nucleotide environment (Figure 1C). We then used the following features to train a generalized linear model (GLM) for the prediction of snoRNA-target interactions:

1. Energy of interaction between the snoRNA and the target, calculated with the PLEXY software for snoRNA target prediction (22).
2. Target site accessibility calculated with CONTRAfold (39).
3. Flanks A content calculated for a window of 30 nts around the putative target site as described in Methods.

We trained the model on chimeric reads involving the rRNA28S and then tested it on chimeric reads involving the rRNA18S. We found that the area under the ROC curve was ∽85% and the model could recall 70% of the known interaction sites with 65% precision (Figure 1E,F). To construct a model for the comprehensive prediction of snoRNA target sites from chimeric reads, we used all known sites in the 28S and 18S rRNAs. At a score threshold of 0.15 we obtain good performance in predicting rRNA modification sites, with a Matthews correlation coefficient (MCC) of 0.19, a precision of 0. 75 and recall value of 0.74 (Figure 2B-D). Therefore, we used this threshold to analyze the genome-wide data of putative snoRNA target sites that were supported by chimeric reads from at least 2 experiments.

### Chimeric reads reveal novel C/D box snoRNA target sites within structural RNAs

We applied the derived model to identify snoRNA interactions with structural RNAs, including beyond rRNAs, snoRNAs, tRNAs and the snoRNAs themselves. Among the 2’-O-Me sites in rRNAs that are presumably guided by C/D box snoRNAs, only one, at position G1383 in the 18S rRNA, is “orphan”, meaning that its guide snoRNA is unknown. Our data indicates that this modification is guided by SNORD30. In addition, in spite of snoRNAs having been studied for a long time, the chimeric reads revealed 39 novel interactions, at 34 distinct methylation sites in rRNAs (16 in rRNA 18S, 21 in rRNA28S and 2 in rRNA5.8S). Eleven of these interactions involve snoRNAs that have been so far classified as “orphan” (Supplementary Table 6). As a specific example, we found that a recently uncovered orphan snoRNA with low but broad expression across tissues, snoID_0701 (family unknown) (Jorjani et al. submitted), was represented among the chimeric reads from two experiments, having a low predicted energy of interaction (−28.2 kcal/mol) with rRNA28S, and the predicted position of modification U2756. Similarly, another snoRNA of unknown family (snoID_372) was found in three experiments to interact with rRNA28S (predicted energy of interaction of −24.8 kcal/mol) being predicted to guide 2’-O-methylation at position 4953.

SnRNAs are also known targets of snoRNA-guided 2’-O-methylation. Of the 9 such sites that are known, we were able to recover 4 with high-confidence and another 2 with less support, but still captured among chimeric reads. Additionally, we identified a novel interaction of SNORD23, a snoRNA that is currently considered orphan, with position 64 of the U6 snRNA (Supplementary Table 6).

Although tRNAs are known to undergo extensive modification including 2’-O-methylation, to our knowledge no evidence has so far been provided for snoRNA-guided modification of tRNAs in animals. In yeast, 2’-O-methylation of riboses in tRNA nucleotides is catalyzed by the methyltransferase Trm7p (48). Here we identified 4 high-confidence interactions of snoRNAs with tRNAs (Supplementary Table 6). Among these, the interaction of the orphan SNORD124 with tRNA Ser-TCY has a relatively low interaction energy of −20.0 kcal/mol and is supported by 3 experiments.

Additionally, we identified 3 interactions of snoRNAs with other snoRNAs. We found an interaction of SNORD5 with SNORD56, of SNORD50 with SNORD57 and of SNORD34 with SNORD38A. One snoRNA, SNORD4B, was found in a chimeric read as both guide and target fragment. Whether all these interactions do indeed lead to 2’-O-methylation or they serve different function remains to be determined (more on this issue below).

### Chimeras capture novel interaction sites outside of typical snoRNA targets

Apart from known and novel snoRNA interactions with structural RNAs the chimeric read data revealed 668 high confidence interactions that did not reside in the typical snoRNA targets, of which 61 (9.1%) involved orphan snoRNAs (Figure 3). The relative frequencies of 2’-O-Me at individual nucleotides were similar to those from known modification sites, where adenines are most frequently targeted (Figure 3A). Based on the ENSEMBL genome annotation we could annotate 433 of the 668 interaction sites, most of which were located in protein-coding genes (303 sites) followed by repeats (45), pseudogenes (18) and lincRNA (12) (Figure 3D). The relative frequencies of targeting different types of genes did not differ substantially between orphan and canonical snoRNAs.

Two of the identified interactions with lincRNAs involved orphan snoRNAs: SNORD73A was found to interact with LINC00954 and snoID_0436 with RP11-554F20.1. The others involved snoRNAs that are known to guide modification of canonical sites in rRNA Supplementary Table 6.

Most of the 303 mRNA-located sites were located in introns (66%), 26% were located in exons and 8% in regions that undergo alternative splicing (annotated as both intronic and exonic, Figure 3E). 35 (45%) of the 78 exonic target sites were located in CDS and 28 (36%) in 3’ UTRs (Figure 3F). These proportion correspond approximately to the total length of different types of regions in the genome, indicating a lack of positional bias of snoRNA binding to mRNAs.

To gain additional insight into the processes that may be regulated by snoRNA binding to mRNAs we analyzed the membership of the mRNAs that were captured in chimeric reads in KEGG pathways as well as their disease associations with the WEB-based GEne SeT AnaLysis Toolkit (http://bioinfo.vanderbilt.edu/webgestalt/) (49). We found that the two most enriched KEGG pathways were glioma and axon guidance (Supplementary Table 7). Correspondingly, the strongest associations were with neurodegenerative diseases and psychiatric disorders including schizophrenia (Supplementary Table 8). For example, SNORD62 appears to target huntingtin (HTT). The consequence of this interaction is however unclear.

SNORD14, SNORD17 and SNORD26 yielded most chimeras with mRNAs (Figure 3B and C). This is not due to their relative abundance, because these snoRNAs are not among the highest expressed. Rather, SNORD14 and SNORD17 were previously reported to have functions beyond 2’-O-methylation of ribose. In particular, SNORD14C was suggested to act as a chaperone in pre-rRNA processing (50) as well as to guide 2’-O-methylation at position 462 of rRNAI 8S. SNORD17 is unusually long for a C/D-box snoRNA (51) and was also identified as one of the small non-coding RNAs that associates with telomeres (52). A role of snoRNAs in alternative processes such as alternative splicing has been suggested before (53) and recently, the interaction of SNORD88 (HBII-180C) with FGFR3 transcripts has been found to influence their splicing (54). Our data contained a chimeric read of SNORD88 with an FGFR3 intron, an interaction predicted to have relatively low energy (−14.2 kcal/mol). However, this interaction is observed in only one experiment and we did not consider it as high confidence.

### Identification of snoRNA-guided 2’O-Me sites with RiboMeth-seq

To further characterize the consequence of the snoRNA-target interactions identified from CLIP data, we applied RiboMeth-seq, a method that was recently developed to accurately map 2’-O-Me sites in rRNAs in high-throughput (2). We carried out seven experiments, six using total RNA, which contains both rRNAs - the canonical snoRNA targets - as well as other RNA species, and one using poly(A)+ RNAs, which is thereby strongly enriched in mRNAs. Two of the total RNA samples were prepared from total cell lysate, two from the nuclear fraction and two from the cytoplasmic fraction. We also developed an improved method for identifying 2’-O-Me sites from RiboMeth-seq data (see Materials and Methods section), and thereby identified 172 high-confidence 2’-O-Me sites, 92 of which novel. 55 (60%) of the novel sites were located in canonical snoRNA targets - snRNA, rRNAs and tRNAs, and only 29 in mRNAs.

From the 753 high-confidence snoRNA target sites identified with CLIP we found that 140, almost exclusively located in rRNAs, are indeed followed by 2’-O-ribose methylation detectable with RiboMeth-seq. These interactions involved 93 distinct target sites, as multiple snoRNAs appear able to interact with the same target site. 117 interactions, at 78 2’-O-Me sites, were already known from previous studies and included 29 out of the 45 known 2’-O-Me sites in rRNA 18S (65%), 46 out of 60 2’-O-Me sites in rRNA 28S (76%), the known site at position 75 in rRNA 5.8S and 2 sites in the U6 snRNA. Figure 4 shows the location of previously known 2’-O-methylation sites in the 18S and 28S rRNAs, as well as the corresponding chimeric read and RiboMeth-seq evidence that we obtained here for these rRNAs.

**Figure 4.**
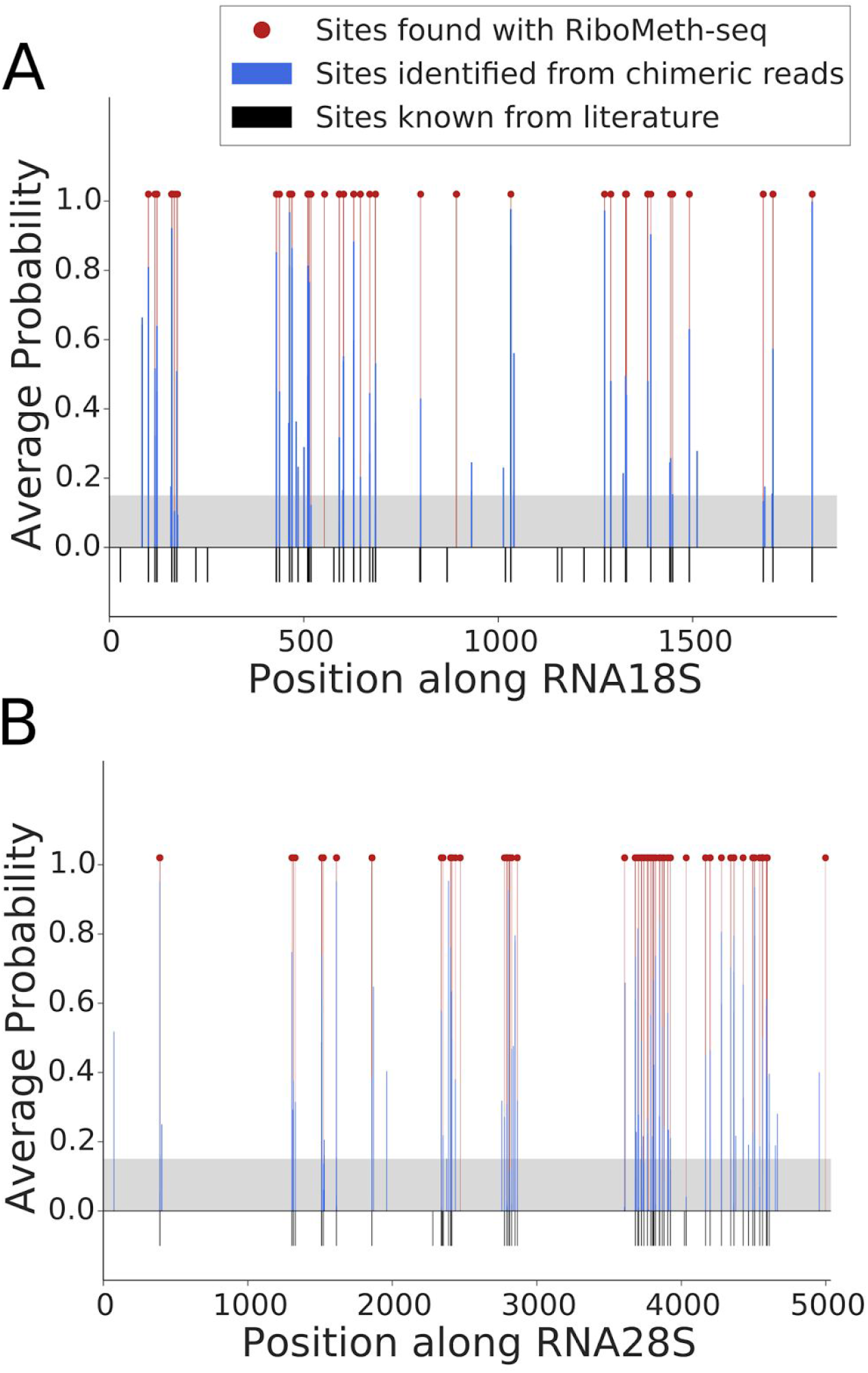
Location of snoRNA interaction sites and 2’-O-ribose méthylation in the 18S (A) and 28S (B) ribosomal subunits. 2’-O-Me positions that are known from literature are shown as black bars. Interactions identified from chimeric reads are shown as blue bars, with their associated probabilities. The grey area indicates the score threshold that we used to extract the high-confidence sites from chimeric reads. The locations of 2’-O-Me sites identified with RiboMeth-seq are shown with red lines and dots.

There are only few novel snoRNA-target interaction sites identified with high confidence from chimeric reads that seem to direct 2’-O-methylation. For example, SNORD2 appears to guide 2’-O-methylation at position G2435 in the 28S rRNA. Almost all others may be due to homology with known 2’O-methylation sites: they were located in different snRNAs isoforms, 2 were in snoRNAs (including an interaction of SNORD4B with itself) and one was in the MTRNR2L6 (Mitochondrially Encoded 16S RNA like 6 protein) transcript, which is similar to the mitochondrially encoded 16S RNA. Thus, the RiboMeth-seq data does not support a role of snoRNAs in guiding 2’-O-methylation at sites with which they interact outside of the structural RNAs.

### Identification of snoRNA guides for novel 2’-O-Me sites obtained with RiboMeth-seq

As mentioned above, 92 of the RiboMeth-seq-derived sites of 2’-O-methylation were novel and also did not correspond to high-confidence sites inferred from chimeric reads. However, not all 2’-O-Me sites are captured reproducibly in chimeric reads and thus, we took a more inclusive approach in assigning snoRNA guides to RiboMeth-seq-derived 2’-O-Me sites. First, we relaxed the constraint that sites should be captured with high confidence in chimeric reads, taking into consideration even single chimeric reads as long as the predicted energy of interaction between the snoRNA and the putative target site was not higher than −6 kcal/mol. With this approach we were able to assign guides to another 5 RiboMeth-seq-supported 2’-O-Me sites in rRNAs: 2’-O-Me of G2468 in the 28S rRNA seems to be guided by two orphan snoRNAs from the snoU13 family (see Jorjani et al. submitted), that of G3606 by SNORD12, of G3771 by SNORD21 and of G4996 by SNORD96B, whereas 2’O-ribose methylation of C174 in the 18S rRNA is guided by SNORD45C.

### Many “orphan” snoRNAs redundantly target known sites of 2’-O-ribose methylation

One of the main open questions in the snoRNA field concerns the targets and functions of the many orphan snoRNAs. At this point, there are 330 orphan snoRNAs that can be grouped into 219 families (Jorjani et al. submitted). Here we found that 66 of these orphan snoRNAs still seem to have canonical targets, acting almost exclusively on known modification sites, which have been previously assigned to other snoRNAs. Among these snoRNAs are:

- SNORD118 - guiding modification of G1612 of rRNA28S (supported by chimeric reads from 6 experiments), as does SNORD80
- snoRNAs of the SNORD115 family - appear able to guide modification of A4560 of rRNA28S, previously assigned to SNORD119. One of these interactions has a very low predicted energy (−20 kcal/mol) and is supported by chimeric reads from 5 experiments
- the recently predicted snoRNA snoID_0719 (Jorjani et al. submitted) - guides the modification of C3680 of rRNA28S, previously attributed to SNORD88
- snoRNAs of the SNORD114 family appear to guide methylation of U4590 of rRNA28S, previously assigned to SNORD72

All the sites are summarized in Supplementary Table 6.

### Assignment of high-confidence 2’-O-Me sites to snoRNAs with PLEXY

Finally, we considered the possibility that no chimeric reads were captured for some of the novel high confidence 2’-O-Me sites identified with RiboMeth-seq. To predict snoRNA guides for these sites, we extracted sequences of 15 nucleotides around the 92 2’-O-Me RiboMeth-seq-based identified sites and carried out snoRNA target predictions with PLEXY, discarding the two sites that were located in PATCH chromosomes. With this procedure we were able to assign snoRNA guides to 54 out of 90 novel methylation sites, leaving 40% of the novel RiboMeth-seq sites without an assigned snoRNA. To many 2’-O-Me site we could assign more than one snoRNA including small non-coding RNAs classified as SNORD-like-snoRNA (Jorjani et al. submitted). This predictions are summarized in Supplementary Table 9. Motif analysis with MEME (55) of the sequences around 2’-O-Me sites that were still left without a snoRNA guide did not reveal any significant motif, and thus it does not appear that they can be assigned to a small number of regulators. Analysis of these regions surrounding these sites with RNAfold (56) did not reveal any significant structure that could explain the capture of these sites in RiboMeth-seq although methylated position tended to be unpaired in comparison to same sequences shuffled (Supplementary Figure 4A and B).

### Evidence of 2’-O-methylation of CLIP-based identified sites in RiM-seq data

Lastly, we wondered whether some of the sites that we captured in chimeric reads were false negatives of the RiboMeth-seq method. We therefore asked whether evidence for their 2’-O-methylation can be found in the RiM-seq data that we obtained in a previous study (Jorjani et al. submitted). Starting from high-confidence RiM-seq-defined sites of 2’-O-methylation we were able to find chimeric read support for chr12:G49050514 (hg19 version of the human assembly from the University of California Santa Cruz, genome.cse.ucsc.edu), which is located in the SNORA2A snoRNA and whose modification seems to be guided by SNORD118, and chr12:G6690685, located in SCARNA11, whose modification could be guided by SNORD6. With this approach we also confirmed two novel RiboMeth-seq and CLIP-supported 2’-O-methylation sites in the 28S rRNA: G3606, whose modification is guided by SNORD12 and G3771, bound by SNORD21. All the results can be found in Supplementary Table 6.

### Low-throughput experimental validation of 2’-O-Me sites

For additional validation of novel 2’-O-Me sites that we identified as described above, we applied the recently published Reverse Transcription at Low deoxy-ribonucleoside triphosphate (dNTP) concentrations, which we then followed with qPCR, to improve quantification (RTL-P (46)). We first tested RTL-P on a known modification site, position A1031 in the human rRNA18S, and obtained a positive result (Supplementary Figure 5A and B). We also performed RTL-P on the site U1991 in 28S rRNA which has no known nucleotide modification (Supplementary Figure 5C). As expected, we obtained a negative result.

**Figure 5.**
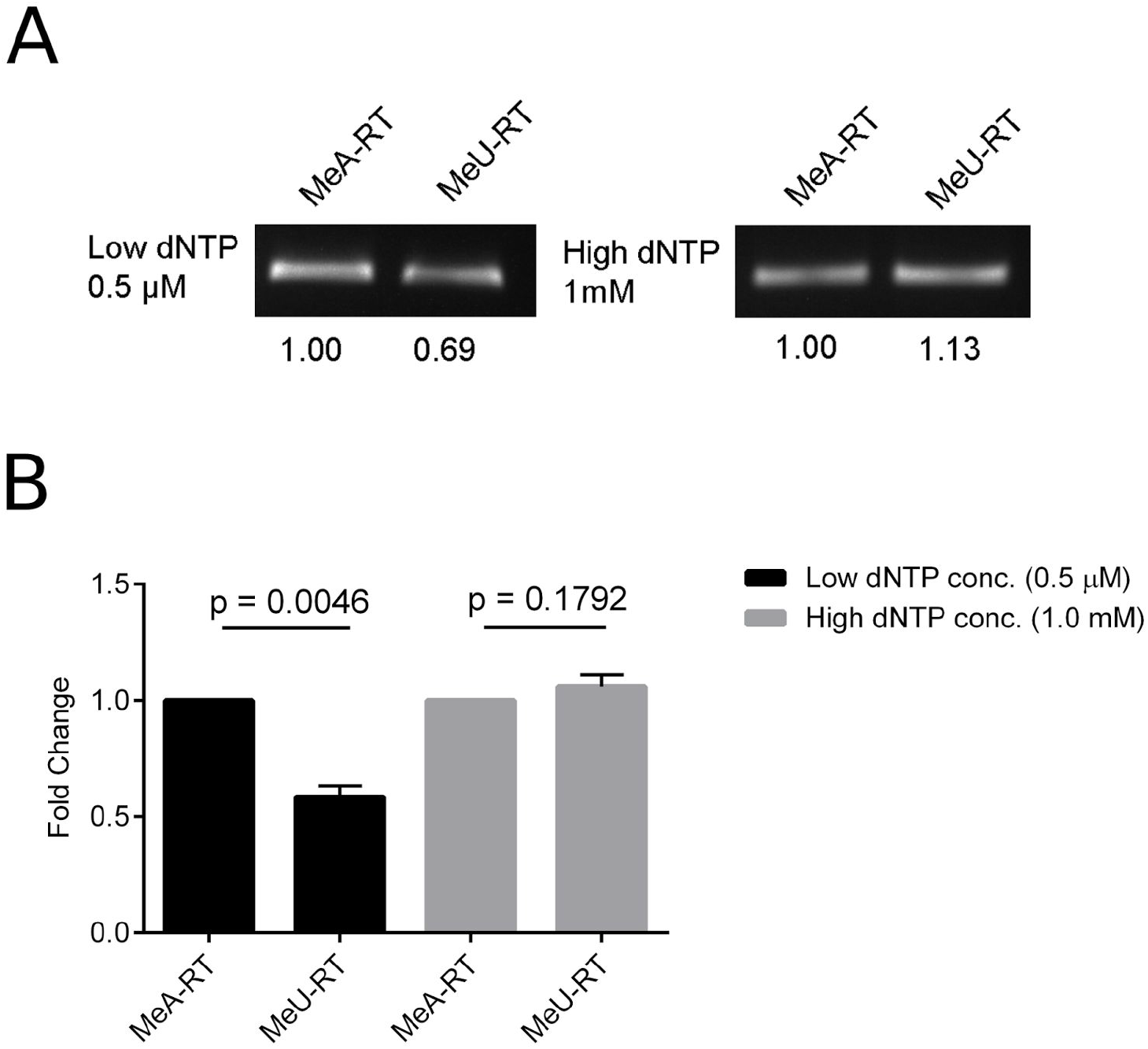
Confirmation of the Gm2435 2’-O-methylation in 28S rRNA by RTL-P. (A) Detection of Gm2435 by RTL-P followed by agarose gel analysis. (B) Detection of Gm2435 by RTL-P followed by qPCR analysis.

Next, we applied the method to position G2435 in rRNA28S, which we identified as a novel 2’-O-Me site based on chimeric reads and RiboMeth-seq. As shown in Figure 5A and B we found that the unanchored MeU-RT primer yielded significantly less cDNA and hence PCR product than the anchored MeA-RT primer at low dNTP concentrations. Although the effect is less pronounced compared to the site A1031 in rRNA18S, there is a significant difference in the PCR products obtained with the two primers at low dNTP concentrations, whereas no such difference can be observed at high dNTP concentrations. This result indicates that sites that emerged from both CLIP as well as RiboMeth-seq are indeed subject to 2’-O-ribose methylation.

### Limited evidence for non-canonical binding of C/D-box snoRNAs

Finally, to allow for the possibility that some of the chimeric reads captured non-canonical snoRNA-target interactions that involve snoRNA regions outside of anti-sense boxes, which will not be identified with PLEXY, we used a general RNA-RNA interaction model implemented in the RNAduplex tool (57) to predict the energy of interaction between the snoRNA and target fragments of chimeric reads (Supplementary Figure 6). Interestingly, two snoRNAs for which we predicted the highest number of non-canonical interactions, SNORD14 and SNORD17, were previously reported to facilitate rRNA processing, participating in the cleavage of the 5’ external transcribed spacer rather than in 2’-O-methylation (50). For SNORD17 the chimeric reads indeed captured a specific snoRNA subsequence that was not one of the anti-sense boxes but rather the sequence AGCCTCAGTTCCTG located at positions 58 to 71 of the snoRNA (between the D’ and C’ boxes). This sequence was also predicted to base-pair stably with target regions (Supplementary Figure 6C and 7A and B). For other snoRNAs the predicted pattern of base-pairing with the target fragments from chimeric reads was diffuse along the snoRNA (eg. SupplementaryFigure 6D). Whether C/D-box snoRNAs carry out functions beyond 2’-O-methylation of ribose or the snoRNAs can be captured in transient interactions that lack functional relevance with molecules such as mRNAs will require additional work and should be addressed by future studies.

## DISCUSSION

High-throughput sequencing technologies coupled with specific treatments and sample preparation have enabled the characterization of transcriptomes at ever increasing depth and resolution. This lead to the realization that the non-coding transcriptome is as large as the protein-coding fraction (58) and to the discovery of new members of all classes of RNAs, including miRNAs and snoRNA (59, 60). The large number of novel molecular species that were discovered increased the need for functional characterization methods, ideally in high-throughput. The aim of our study was to provide such methods for a specific class of non-coding RNAs, the C/D-box snoRNAs. A recent study to which we contributed expanded the catalog of human snoRNAs by 41 CD-box new molecules (Jorjani et al. submitted) and tried to identified snoRNA targets for the 118 C/D box snoRNAs that were up to that point considered “orphan”.

Here we have combined two high-throughput approaches, the first of which aims to identify direct interactions between snoRNAs and targets and the other to map sites of 2’-O-methylation genome-wide. Because many small RNAs function as guides for ribonucleoprotein complexes, the mapping of RNA-RNA interactions has initially been done indirectly, by crosslinking and immunoprecipitation of proteins from these complexes (28, 29, 61). Due to the relatively weak sequence complementarity constraints and because the small RNA guides typically form families of sequence-related molecules, inferring the small RNA that guided the interaction with each presumed target site determined by CLIP has not been trivial. This led to efforts to capture RNA-RNA interactions directly, which has been accomplished initially through the addition of an RNA-RNA ligation step in CLIP (25, 26). Very recently, through careful analysis of CLIP data, Grosswendt and colleagues found that a specific ligation is not necessary, because endogenous ligases already generate miRNA-target site chimeras that can be found in CLIP samples (30). Whether this is also the case for CLIP carried out with the core snoRNP proteins has not been investigated. Developing a computational data analysis method for the identification of C/D box snoRNA-target hybrids from Nop56/Nop58/Fibrillarin-CLIP data was the first aim of our study. Due to the relatively short length of the snoRNA and target fragments that are captured with CLIP, the relative rarity of chimeric sequences, in the range of less than a percent of the reads (30) and the large proportion of “background” sequences in CLIP (29), a careful analysis that considers the specific base-pairing pattern of snoRNAs with targets is necessary. Using the sites in rRNA28S we have trained a model that shows very good sensitivity and specificity in predicting the 2’-O-Me sites in the rRNA18S. We applied this model to all chimeric sequences from 7 CLIP experiments and identified 753 high-confidence snoRNA-interaction sites including 39 novel interactions in rRNAs.

To investigate the consequences of the identified interactions, we have adapted a high-throughput method, RiboMeth-seq, that was previously applied to the identification of 2’-O-Me sites in yeast rRNAs (2) and we have developed an improved method for the computational analysis of these data. We found that the approach identifies 72% of the known 2’-O-Me sites in rRNAs with a precision of 85%. Interestingly, we found that many of the snoRNAs that were considered “orphan” seem to guide interactions with sites that have already been assigned to other snoRNAs. That is, it appears that there is substantial redundancy of snoRNAs in guiding specific modifications, whose importance for cellular function deserves further investigation. Combining the CLIP with RiboMeth-seq we have validated with high confidence 104 of the 190 known snoRNA-guided interactions with structural RNAs as well as their consequence, namely the 2’-O-methylation of the target. Only 14 of the known 2’-O-Me sites could not be validated (Supplementary Table 10). These results indicate that our approach is very well suited for the characterization of C/D box snoRNA-target interactions in high-throughput. Additional validation of a novel rRNA modification site with an independent, low throughput method ((46)) indicates that the CLIP-RiboMeth-seq approach is reliable. Although in species such as yeast or human these interactions have been extensively studied for a relatively long time and much is already known, the approach should be suitable for other species as well.

Our analysis indicates also that neither of the methods is fully comprehensive in its coverage. Although this could be due to some extent to the cell type-specificity of expression or inter-molecular interactions, it is the case that guide RNA-target chimera are captured with low efficiency with the approaches that are currently available (26, 30) and in this aspect, there is room for further improvement. In RiboMeth-seq, a low expression of the 2’-O-methylated RNAs could hinder their identification. This could be a problem for the identification of modification sites in targets that are expressed in a tissue-specific manner, in mRNAs or lincRNAs, whose abundance per cell is much lower compared to the rRNAs. This could be a reason for the lack of experimental validation of 2’-O-methylation of mRNA target sites that emerged from chimeric reads. The data that we obtained with the RTL-P approach also suggests that at least some of the novel sites of snoRNA-target interaction are not methylated with 100% efficiency, indicating that an increased sensitivity of detection of 2’-O-Me sites may confirm the role of newly discovered interactions in 2’-O-ribose methylation. Alternatively, not all snoRNA-target interactions lead to 2’-O-methylation. Indeed, an ancestral function of snoRNAs in rRNA processing that is still preserved in the U3 snoRNA has been proposed (20). Which snoRNAs have retained an ancestral function or have acquired functions beyond 2’-O-methylation remains to be determined. RiboMeth-seq also revealed a few high-confidence sites for which we did not find any corresponding chimeric reads. Although this could be due to the limited coverage of snoRNA-target interactions by chimeric reads, they could represent sites that are resistant to alkaline hydrolysis for reasons other than 2’-O-Me. Supporting this latter hypothesis, these sites are generally located in rRNAs or snRNAs, molecules that are extensively modified and highly structured. Moreover, in contrast to rRNA-known modification sites, which do not exhibit any nucleotide bias, the new sites recovered by RiboMeth-seq show a strong G-bias (Supplementary Figure 3F). This could indicate that these modifications are introduced by specific enzymes such as the transfer RNA methyltransferase 7 protein (62).

Our approach would be particularly interesting to apply to study the role of C/D box snoRNAs in development or across cell types. Indeed, in updating the catalog of human snoRNAs, we have found that a substantial proportion of snoRNAs has a strong specificity for neuronal cells, and that there is a snoRNA expression signature of cancers (Jorjani et al. submitted). Interestingly, although we performed our analysis in HEK cells, analysis of disease associations of the mRNAs that were captured in chimeric reads revealed that most targeted diseases are neurological disorders. The significance of snoRNA expression changes can be studied with the techniques we established here.

## ACKNOWLEDGEMENTS

R.G. and D.J. would like to thank the members of the Zavolan lab for input on the project.

